# A small viral potassium ion channel with an inherent inward rectification

**DOI:** 10.1101/559716

**Authors:** Denise Eckert, Tobias Schulze, Julian Stahl, Oliver Rauh, James L Van Etten, Brigitte Hertel, Indra Schroeder, Anna Moroni, Gerhard Thiel

## Abstract

Some algal viruses have coding sequences for K^+^ channels with structural and functional characteristics of pore modules of complex K^+^ channels. Here we exploit the immense structural diversity of natural channel orthologs to discover new basic principles of structure/function correlates in K^+^ channels. The comparative analysis of three similar K^+^ channels with monomer sizes ≤ 86 amino acids (AA) shows that one channel (Kmpv_1_) generates an ohmic conductance in HEK293 cells while the other two channels (Kmpv_SP1_, Kmpv_PL1_) exhibit typical features of canonical Kir channels. Like Kir channels, the rectification of the viral channels is a function of the K^+^ driving force. Reconstitution of Kmpv_SP1_ and Kmpv_PL1_ in planar lipid bilayers showed rapid channel fluctuations only at voltages negative of the K^+^ reversal voltage. This rectification was maintained in KCl buffer with 1 mM EDTA, which excludes blocking cations as the source of rectification. This means that rectification of the viral channels must be, unlike Kir channels, an inherent property of the channel proteins. The structural basis for rectification was investigated by a chimera between rectifying and non-rectifying channels as well as point mutations, which made the rectifying channels similar to the ohmic conducting channel. The results of these experiments exclude the domain, which connects the two transmembrane helixes and which includes the pore helix and the selectivity filter, as playing a major role in rectification; inward rectification must be conferred by the transmembrane domains. The finding that a swapping of the AA, which is typical for the two inward rectifiers, with the respective AA from Kmpv_1_ did not compromise rectification suggests that tertiary or quaternary structural interactions are responsible for this type of gating.

## INTRODUCTION

The key properties of K^+^ channels such as ion selectivity and gating are well understood based on high-resolution structures (Doyle et al., 1998; Perozo, 2002; MacKinnon, 2003; Kuang et al., 2015; Roux, 2017; Nichols and Lee, 2018). However, despite this progress many functional aspects and in particular the significance of individual amino acids (AA) for function cannot be easily obtained from static protein structures. Therefore, other approaches are still needed to understand structure/function correlates in K^+^ channels. In recent years “assumption free” genetic methods in combination with functional assays were developed as powerful tools for uncovering functional alterations, which are caused by distinct mutational changes (Minor, 2009).

In this context the small K^+^ channel proteins from viruses turn out to be an interesting model system for understanding functional aspects in the pore module of K^+^ channels. From a structural and functional point of view the viral channels represent the “pore module”, which is present in all known K^+^ channels (Thiel et al., 2011). Because of this common architecture any insights from the viral channels are probably relevant for the function of complex human K^+^ channels (Gazzarrini et al., 2003). A further advantage of the viral channels as a model system is that they are the smallest proteins known to form a functional K^+^ channel. This combination of small size and robust function limits the complexity of the system to less than 100 AAs.

Previous experiments have established that the activity of some viral K^+^ channels is essential for infection of their hosts (Thiel et al., 2010). The implication is that the viral genes are under evolutionary pressure and that their gene products need to form functional channels (Hamacher et al., 2012). This assumption has been supported by experimental data, which have shown that the AA sequences of viral K^+^ channels are variable and that the gene products are still functional in different test systems (Kang et al., 2004; Gazzarrini et al., 2004; Rauh et al., 2017). The sequence variability of viral K^+^ channels, which can be isolated from various environmental samples, results in a large library of variable K^+^ channel sequences with functional variability. We have exploited this structural diversity and have identified interesting functional differences, which are rooted in the sequence variability in these channels. The power of this unbiased approach is best illustrated by the fact that even very conservative AA exchanges caused significant functional differences. In the Kcv channel from chloroviruses; e.g., an exchange of Phe for Val or Leu for Iso in the first transmembrane (TM) domain drastically altered the Cs^+^ sensitivity of the channel as well as its voltage dependency (Kang et al., 2004: Gazzarrini et al. 2004). The results of these experiments underscored the importance of the outer TM domain for K^+^ channel function, which had largely been ignored. In the Kcv channel scaffold from SAG chloroviruses it was found that a mutation of Gly versus Ser in the inner transmembrane helix (TM1) affected the open probability of the channel. A closer investigation of these mutations uncovered a new type of gating mechanism, which is based on an intra helical hydrogen bond between the critical Ser and an upstream partner AA in the alpha helix (Rauh et al., 2017).

Here we further exploit the diversity of viral K^+^ channel genes by screening viral K^+^ channels from a marine habitat. We have previously reported that some viruses, which infect unicellular marine algae, also encode genes with the hallmarks of K^+^ channels (Siotto et al., 2014). An initial functional testing of some of these proteins revealed that they have non-canonical architectures in their TM domains, but that they still form functional K^+^ channels (Siotto et al., 2014). Here we perform a comparative examination of K^+^ channels, which are similar in their structure but fundamentally different in their voltage dependency. While one channel generates an ohmic conductance the other two proteins exhibit a typical Kir-like inward rectification in which large inward currents occur only at membrane voltages negative to the K^+^ equilibrium potential. The data show that this rectification is an inherent property of the protein and does not require Mg^2+^ or polyamines as a blocker. By mutational studies we identify the N-terminal part of the outer TM domain as a crucial element for an inherent inward rectification of this channel.

## MATERIALS AND METHODS

The electrical properties of the putative viral channels in HEK293 cells were recorded as reported previously (Siotto et al., 2014). Constructs of Kmpv_1_ and Kmpv_SP1_ were transiently expressed as fusion proteins with GFP on the C-terminus using the liposomal transfection reagent GeneJuice^®^ (MERCK KGaA, Darmstadt, Germany). Measurements were performed at room temperature in a bath solution containing: 1.8 mM CaCl_2_, 1 mM MgCl_2_ and 5 mM 4-(2-hydroxyethyl)-1-piperazineethanesulfonic acid (HEPES, pH 7.4) and either 50 mM KCl or 50 mM NaCl; different concentrations of BaCl_2_ were added to the K^+^ containing media to block the channels. The pipette solution contained 130 mM potassium-D-gluconic acid, 10 mM NaCl, 5 mM HEPES, 0.1 mM guanosine triphosphate (Na salt), 0.1 μM CaCl_2_, 2 mM MgCl_2_, 5 mM phosphocreatine and 2 mM adenosine triphosphate (Na salt, pH 7.4). The osmolarity of all solutions was adjusted with mannitol to 330 mosmol/kg. For the standard solutions we used JPCal software (Barry, 1994) to calculate a liquid junction potential of 15 mV, which was subtracted from the clamp voltages.

Planar lipid bilayer experiments were done with a vertical bilayer set up (IonoVation, Osnabrück, Germany) as described previously (Braun et al., 2014). A 1% hexadecane solution (MERCK KGaA, Darmstadt, Germany) in n-hexane (Carl ROTH, Karlsruhe, Germany) was used for pretreating the Teflon foil (Goodfellow GmbH, Hamburg, Germany). The hexadecane solution (ca. 0.5 μl) was added in the hole (100 μm in diameter) in the Teflon foil with a bent Hamilton syringe (Hamilton Company, Reno, Nevada, USA). The experimental solution contained 100 mM KCl and was buffered to pH 7.0 with 10 mM HEPES/KOH. As a lipid we used 1,2-diphythanoyl-sn-glycero-3-phosphocholine (DPhPC) (Avanti Polar Lipids, Alabaster, AL, USA) at a concentration of 0.15-25 mg/ml in n-pentane (MERCK KGaA, Darmstadt, Germany).

The Kmpv_PL1_, Kmpv_SP1_ and Kmpv_1_ proteins were synthetized cell-free, purified over a Ni-NTA column and reconstituted into planar lipid bilayers directly as described (Winterstein et al., 2018).

Data are generally presented as mean ± standard deviation (sd) of n independent experiments. Statistical significance was evaluated by student T-test.

## RESULTS

### Similar channels with different functional properties

We screened a library of small and structurally similar K^+^ channels from marine phycodnaviruses (Siotto et al., 2014) for interesting functional differences. In this search we found three channels with similar lengths and high AA identity. Two of the channels, Kmpv_PL1_ and Kmpv_SP1_, are 75% identical; both channels differ from the third channel Kmpv_1_ in 29 AAs of which 16 are conservative or semi-conservative AA exchanges (Figure 1A). In spite of the structural similarities a comparative functional analysis of Kmpv_1_ and Kmpv_SP1_ in HEK293 cells uncovered striking functional differences with respect to gating and Ba^2+^ sensitivity.

**Figure 1:**
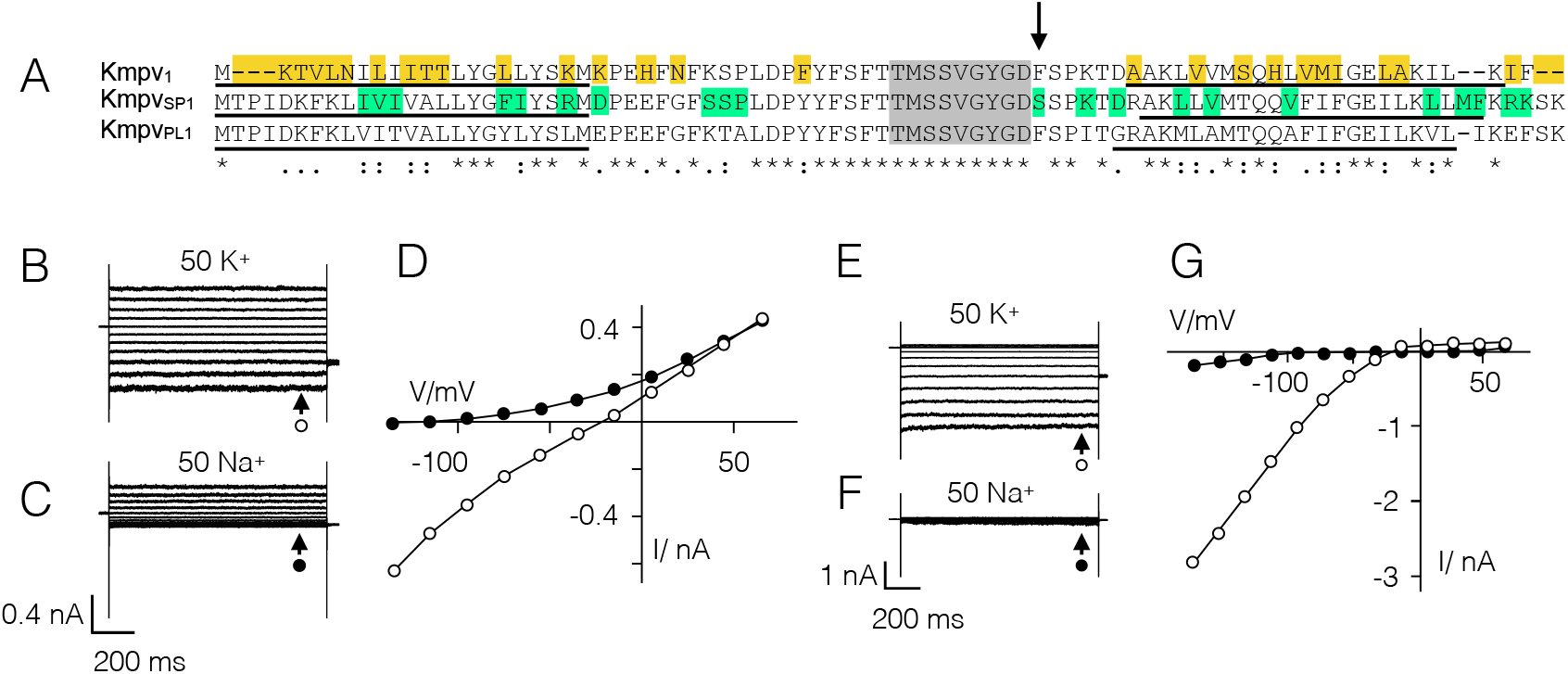
Similar channels with partially different functional properties. **(A)** Alignment of three channel proteins Kmpv_1_, Kmpv_PL1_ and Kmpv_SP1_. The identical filter domain is highlighted in grey. The amino acids in Kmpv_1_, which are different from the two other proteins, are indicated in yellow. The amino acids in which Kmpv_PL1_ differs from Kmpv_SP1_ are marked in green. The estimated transmembrane domains are underlined. A crucial AA 52 in Kmpv_SP1_ is marked by an arrow. Exemplar current responses to voltage steps between +65 mV and -135 mV of HEK293 cells expressing Kmpv_1_ in 50 mM K^+^ **(B)** and 50 mM Na^+^ **(C)**. The corresponding steady state I/V relations are shown in **(D)**. Same experiments with cell expressing Kmpv_SP1_ in 50 mM K^+^ **(E)** and 50 mM Na^+^ **(F)** plus corresponding I/V relations in **(G)**. In both I/V relations open symbols report measurements in K^+^ and closed symbols report measurements in Na^+^. Note that Kmpv_1_ generates an ohmic conductance in K^+^ and Kmpv_SP1_ exhibits a strong inward rectification. Asterisks (*) indicate conserved residues, colons (:) indicate residues with very similar properties, and periods (.) indicate residues with weakly similar properties.

The difference in gating is apparent in Fig. 1 and Fig. 2: while Kmpv_1_ generates an ohmic conductance in HEK293 cells (Fig. 1B,D), Kmpv_SP1_ produces a strong inward rectification (Fig. 1E,G). Even though both channels exhibit a similar selectivity for K^+^ over Na^+^ (Fig. 1C,F; Table 1) they differ in their sensitivity to the canonical K^+^ channel blocker Ba^2+^ (Fig. 2). While the ohmic Kmpv_1_ channel is moderately sensitive to Ba^2+^ (Fig. 2A,D) the inward rectifier Kmpv_SP1_ exhibits a stronger voltage dependent block (Fig. 2G,H). The dose dependency for a Ba^2+^ block of Kmpv_SP1_ (Fig. 2C,D) has at -115 mV a half maximal inhibition at 14 μM Ba^2+^; Kmpv_1_ requires an ca. 100-fold higher concentration for the same degree of inhibition.

**Figure 2:**
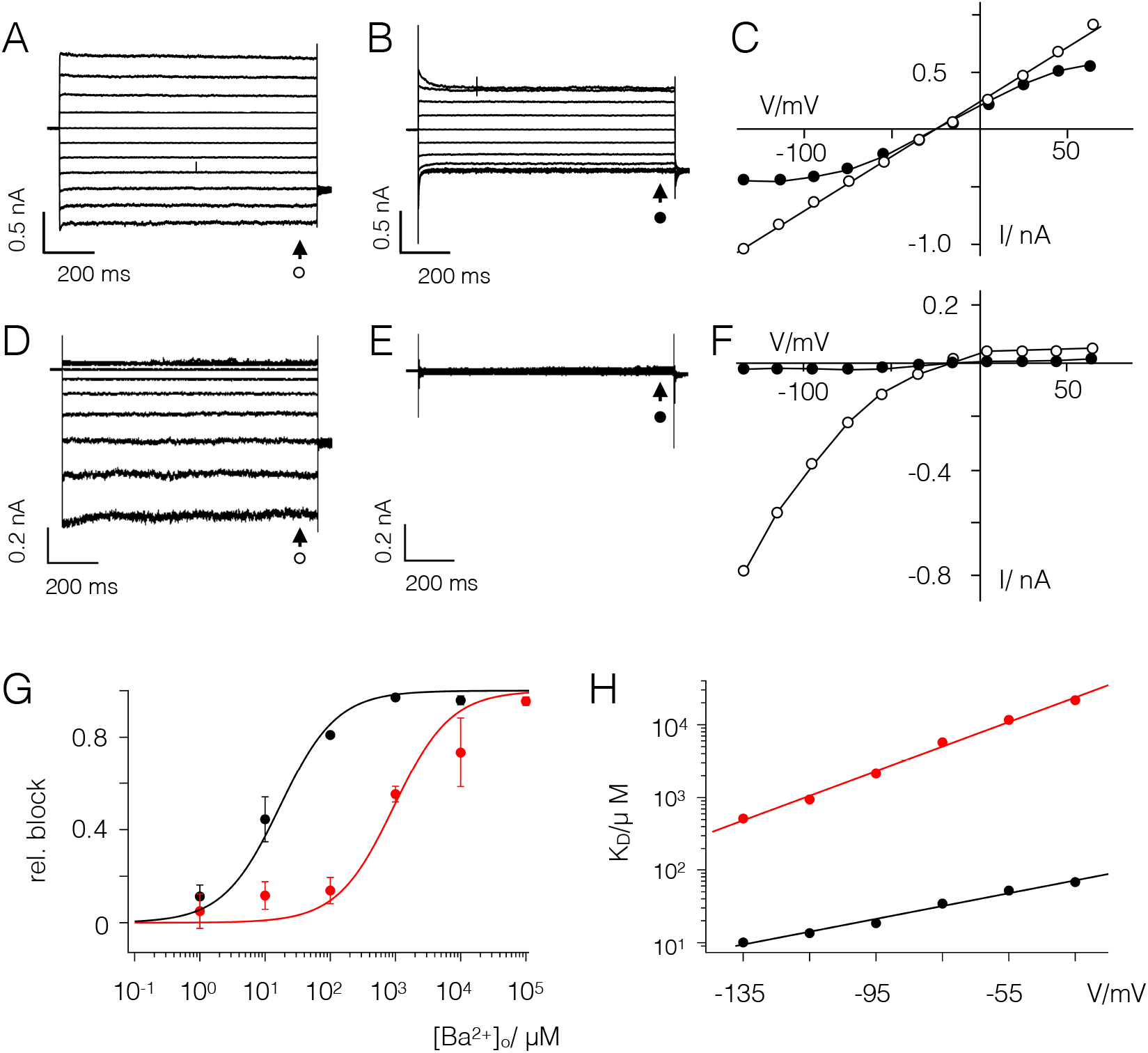
Kmpv_1_ and Kmpv_SP1_ exhibit a different sensitivity to extracellular Ba^2+^. Exemplar currents and corresponding I/V relations as in Fig. 1 from HEK293 cells expressing Kmpv_1_ **(A,D)** or Kmpv_SP1_ **(B,E)**. Currents were recorded in bath solution with 50 mM K^+^ in absence **(A,B)** and presence of 1 mM BaCl_2_ **(D,E)**. The corresponding I/V relations for Kmpv_1_ **(C)** and Kmpv_SP1_ (F) show steady state currents in the absence (open symbols) and presence (closed symbols) of 1 mM Ba^2+^. **(G)** Relative block of Kmpv_SP1_ (black) or Kmpv_1_ (red) at -115 mV as a function of extracellular Ba^2+^ concentration. Data (mean ± sd; n≥4) were fitted with the Hill equation (eqn. 1). The K_D_ values for Ba^2+^ block of both channels from fits as in G are plotted as a function of voltage **(H)**. The K_D_ value increases ten fold per 114 mV (Kmpv_SP1_, black) and 59 mV (Kmpv_1_, red) of negative voltage.

**Table 1.** Kmpv_1_ and Kmpv_SP1_ are moderately selective for K^+^ over Na^+^. Reversal voltages (V_rev_, mean ± sd; number of experiments in brackets) were measured in HEK293 cells expressing either Kmpv_SP1_ or Kmpv_1_ with 130 mM K^+^ and 10 mM Na^+^ in the pipette and either 50 mM K^+^ or 50 mM Na^+^ in the bath medium. The relative permeability ratio P_K+_/P_Na+_ was calculated with Goldman equation based on measured Vrev values (numbers in brackets denote number of independent recordings). For comparison the permeability ratio P_K+_/P_Na+_ ratio of Kir channel ROMK1 is > 100 (Spassova and Lu, 1999).

| Channel | $V_{rev}$ (K<sup>+</sup>) | $V_{rev}$ (Na<sup>+</sup>) | $P_{K^+}/P_{Na^+}$ |
| --- | --- | --- | --- |
| Kmpv<sub>SP1</sub> | -22±3 (55) | -102±4 (7) | 24 |
| Kmpv<sub>1</sub> | -23±2 (13) | -100±7 (8) | 21 |

### The inward rectification is an inherent function of Kmpv_SP1_

The voltage dependency of Kmpv_SP1_ (Figs. 1G, 2F) resembles the strongly inward rectifying Kir channels (Hibino et al., 2010). To test if the voltage dependency in Kmpv_SP1_ is like Kir channels, which depend on the driving force for K^+^ ions, we recorded currents over a range of K^+^ concentrations in the bath. The representative current responses to voltage ramps show that the rectification shifts like in Kir channels with the K^+^ reversal voltage (Fig. 3A). The channel shows no appreciable conductance at voltages positive of the K^+^ reversal voltage (V_K+_). These data suggest that Kmpv_SP1_ functions similar to Kir channels.

**Figure 3:**
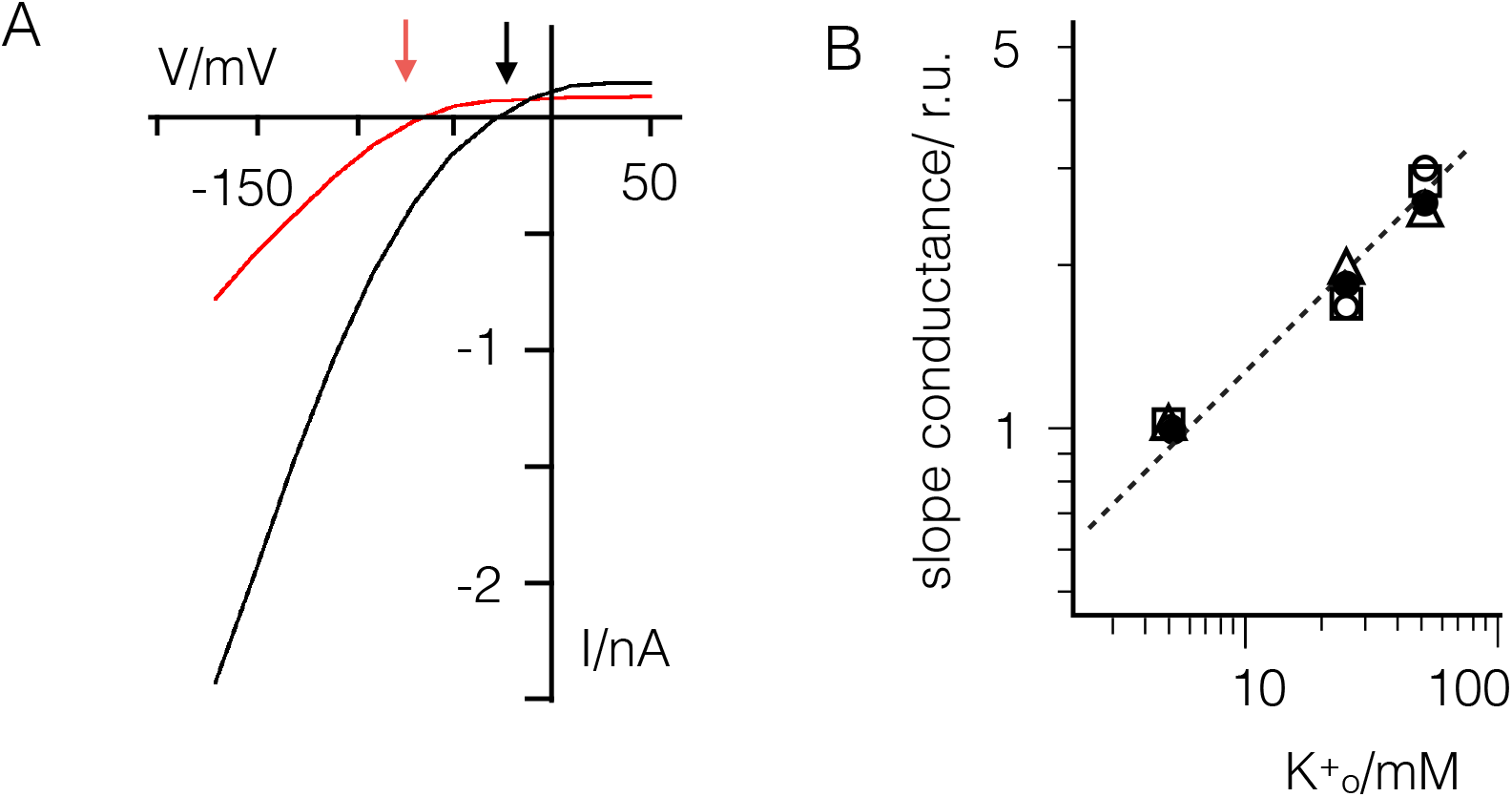
Inward rectification of Kmpv_SP1_ shifts with driving force for K^+^. **(A)** Exemplar current responses of one HEK293 cell expressing Kmpv_SP1_ to voltage ramp between 50 and -160 mV with 5 mM (red) or 50 mM (black) K^+^ in the bath solution. Large currents occur negative of K^+^ reversal voltages, which are indicated by arrows for 5 and 50 mM K^+^. **(B)** Slope conductance of KmpvSP1 generated inward current between -95 and -135 mV as a function of the extracellular K^+^ concentration. Data from n=4 measurements were normalized to conductance in 5 mM K^+^ and jointly fitted with eqn. 1 yielding a value of 0.46 for n.

A hallmark of many Kir channels is that the slope conductance of the inward current has a square root dependency on the extracellular K^+^ concentration (Lopatin and Nichols, 1996; Makhina et al., 1994; Perier et al., 1994). To determine the respective dependency for Kmpv_SP1_ we measured the slope conductance of Kmpv_SP1_ between -135 and -95 mV in solutions with K^+^ concentrations between 5 and 50 mM. The values from 4 measurements are shown in a double logarithmic plot (Fig. 3B). The data can be jointly fitted with equation (1)

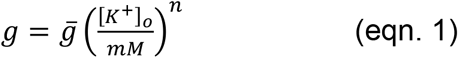

where g is the *slope* conductance, ḡ the reference *slope* conductance (5 mM external K^+^ and constant 130 mM internal K^+^). [*K*^+^]_*o*_ gives the variable external K^+^-concentration and n is a constant. The best fit is obtained with a n value of 0.46 (Fig. 3B), which indicates that the slope conductance is approximately proportional to the square root of the external K^+^ concentration.

The data so far indicate that the viral channel shares some but not all functional features of Kir channels. To examine the mechanism, which allows rectification, Kmpv_SP1_ and Kmpv_PL1_ were synthesized *in vitro* and reconstituted in planar lipid bilayers. For comparison the ohmic Kmpv_1_ channel was examined in the same manner.

The representative data in Fig. 4 show that Kmpv_1_ generated current fluctuations with a small unitary conductance of ca. 6 pS and a high voltage independent open probability. The time averaged I/V relation, e.g., the product of unitary conductance and open probability, is largely linear over a voltage window of ±100 mV. This is consistent with the view that this channel is generating the ohmic I/V relation of Kmpv_1_ measured in HEK293 cells (Fig. 1 B,D).

**Figure 4:**
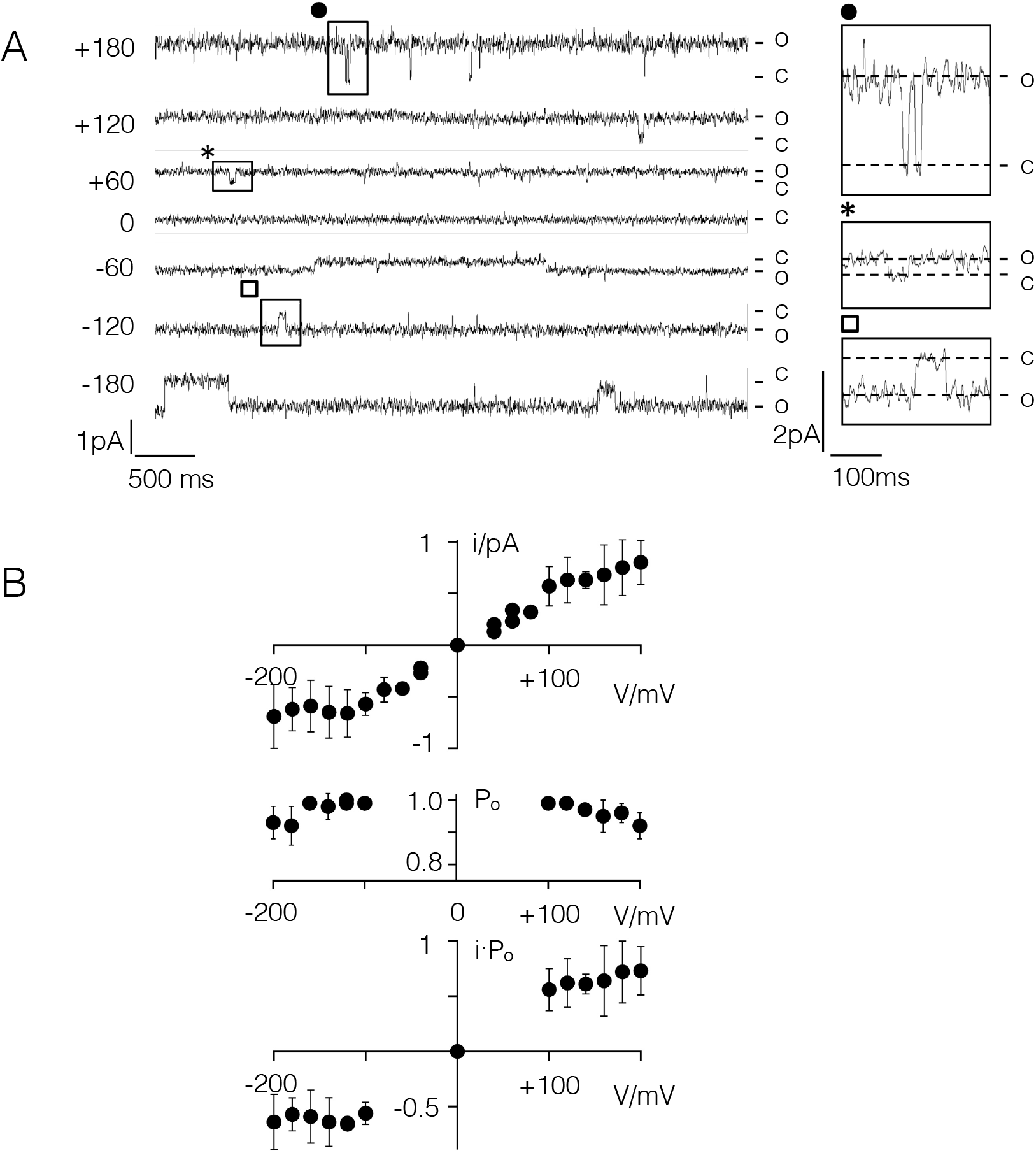
Kmpv_1_ generates a voltage independent channel with small unitary conductance in planar lipid bilayers. **(A)** Exemplar current traces of Kmpv_1_ activity over a range of clamp voltages in symmetrical KCl (100 mM + 10 mM HEPES, pH 7). The channel protein was translated *in vitro* into nanodiscs and recorded in a vertical planar DPhPC bilayer (Winterstein et al., 2018). The channel exhibits a high open probability with only some longer-lasting closures. Individual closing events, which are marked on the traces, are enlarged on the right. **(B)** Mean unitary channel voltage relation (top), open probability voltage relation (middle) and time averaged I/V relation (bottom) from n= 2 to 5 recordings. The latter were obtained by multiplying values from unitary I/V and P_o_/V relations.

The currents generated by Kmpv_SP1_ are different. The representative single channel recordings show that this protein exhibits a high flicker type activity at negative voltages with only some longer lasting resolvable closed times (Fig. 5A). These fluctuations are only evident at negative but not at positive voltages resulting in a strong inward rectifying I/V relation of the mean current (Fig. 5B). The I/V relation from the bilayer measurements is similar to that recorded in HEK293 cells under comparable ionic conditions (Fig. 5C). In contrast to Kir channels, the viral inward rectifier does not require the PIP2 phospholipid for activity (Huang et al., 1998). The data furthermore underscore that the flicker type channel fluctuations in the bilayer are indeed responsible for the Kmpv_SP1_ currents in HEK293 cells. This assumption is further supported by the finding that 1 mM Ba^2+^ greatly reduced the flickering channel activity in bilayers at negative voltages (Fig. 5A,B).

**Figure 5:**
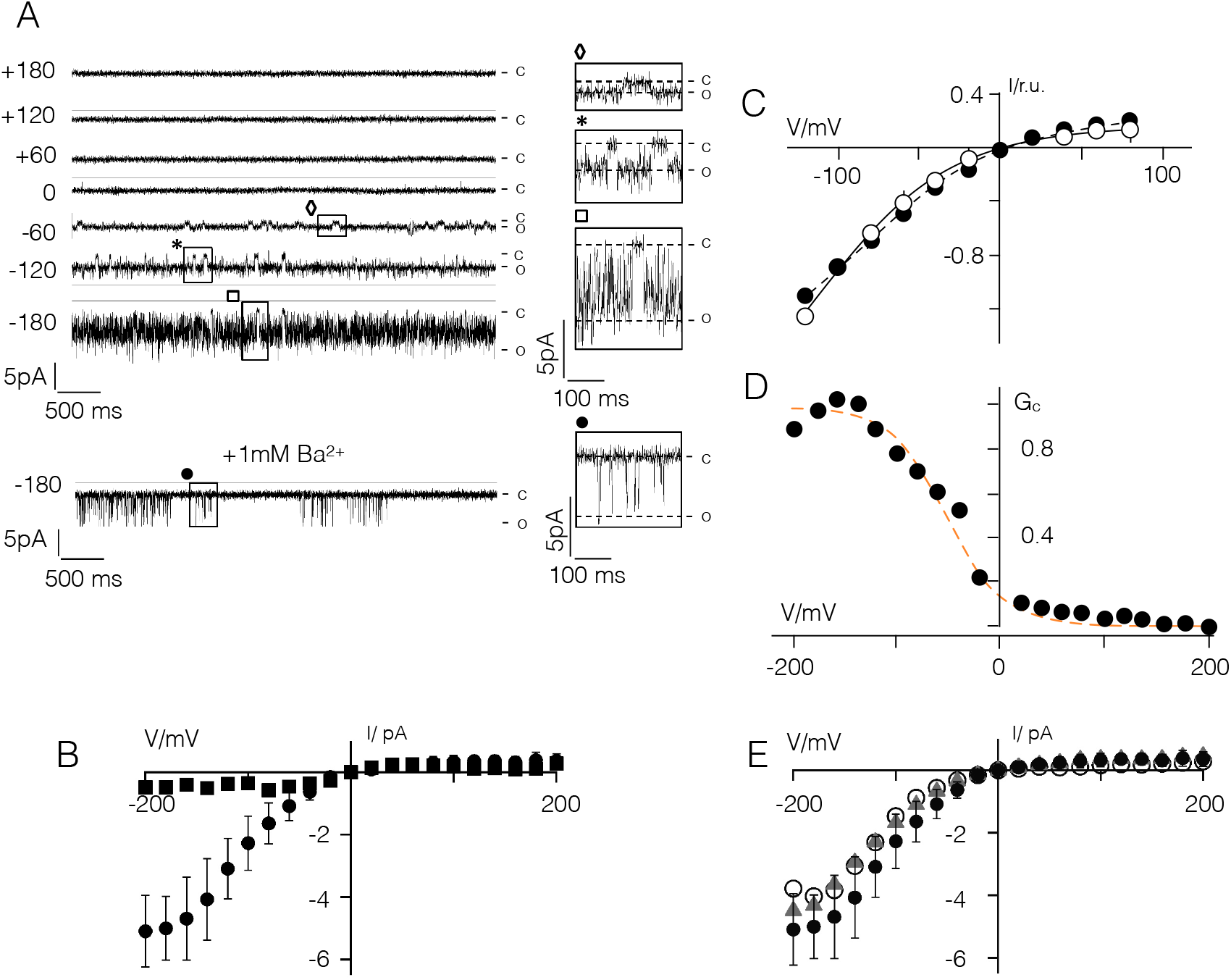
Kmpv_SP1_ has an inherent inward rectification. **(A)** Exemplar current traces of Kmpv_SP1_ activity in bilayer recordings over a range of clamp voltages in symmetrical KCl (100 mM KCl + 10 mM HEPES, pH 7) in absence or presence of BaCl_2_ (+1 mM Ba^2+^). Clamp voltages are given (in mV) on the left of current traces. The channel protein was translated *in vitro* as in Fig. 2. The channel exhibits a high open probability with flicker type fluctuations. Individual clear-cut closing events, which are marked on the traces, are enlarged on the right. **(B)** I/V relation of mean currents measured over clamp steps of 60 sec in absence (circles, mean ± sd; n=9) and presence of Ba^2+^ (squares). **(C)** Normalized mean I/V relations of Kmpv_SP1_ currents (mean ± sd; n≥4) measured in bilayers (open circles) and in HEK293 cells as in Fig. 1 but with 100 mM K^+^ in bath solution (closed circles). **(D)** Chord conductance (Gc)/voltage relation of mean current from B fitted with Boltzmann function (orange line) using z value of =0.9. **(E)** I/V relation of Kmpv_SP1_ channel from bilayer recordings as in B with 10 mM EDTA in *cis* and *trans* chamber (triangles) or after removing HEPES from the bath solution (open circles). For comparison the data from B are re-plotted (closed circles).

To estimate the steepness of the voltage dependency in the Kmpv_SP1_ channel we plotted the relative chord conductance (G_rel_) of the mean current from Fig. 5D as a function of voltage. The data were fitted to a Boltzmann function (dotted line) of the form G = (1 + e^zF(V − V_1/2_)/RT^)^−1^ where z is the effective charge (=charge * electrical distance), V_1/2_ the half activation voltage, F, R and T have their usual thermodynamic meaning. This yields values of -48 mV and 0.9 for V_1/2_ and z, respectively. The results of this analysis indicate that the inherent voltage dependency of the Kmpv_SP1_ channel is more shallow than that induced by spermidine in canonical Kir channels but in the range of the voltage dependency generated by Mg^2+^ block in these channels (Lopatin et al., 1994, Kurata et al., 2004; Lu, 2004).

While expression of Kmpv_PL1_ generated no current in HEK293 cells it was successfully reconstituted in bilayers using the same procedure used for Kmpv_SP1_. The data in Fig. 1 supplement show that this protein generated the same inward rectifying I/V relation as the related Kmpv_SP1_ protein. From these data we conclude that the amino acid differences between these two channels (Fig. 1A) can be excluded as the origin of rectification.

In Kir channels inward rectification is generated by a voltage dependent block by cytosolic Mg^2+^ or polyamines (Baronas and Kurata, 2014; Nichols and Lee, 2018). The finding that Kmpv_SP1_ rectifies in a bath solution with only KCl and HEPES in the same manner as in cells suggests that the channel may harbor an inherent mechanism of rectification. To exclude the possibility of HEPES (Guo and Lu, 2002) or any contamination of divalent cations in the bath solution as a source of rectification, we performed experiments as in Fig. 5A in an un-buffered pure KCl solution. We also measured channel activity in a KCl/HEPES buffer plus 1 mM EDTA in the *cis* and *trans* chamber to chelate any possible divalent ion contaminant. In both cases Kmpv_SP1_ generated the same inward rectifying I/V relation as in the standard recording condition (Fig. 5E). In additional experiments channel activity was measured in a 100 mM KCl/HEPES buffer plus 1 mM EDTA (in the same buffer as before) for more than 0.5 h. We reasoned that the affinity of a binding site in the channel to a putative blocker might be very high and that it might take time to dissociate (Kurata et al., 2011). However, when we compared the I/V relations from measurements at the beginning of the experiments with those recorded 0.5 h later, we found no difference; such a long incubation time is sufficient to release blockers from the channel pore of canonical Kir channels (Lopatin et al., 1994) (Fig. 2 supplement). Collectively these data suggest that Kmpv_SP1_ has an inherent mechanism of gating, which generates a phenotype similar to that of Kir channels.

### Structural basis of rectification

After finding functional similarities and differences between the viral inward rectifiers and canonical Kir channels we aligned Kmpv_SP1_ with representatives (Kir1.1, Kir2.1, Kir3.1 and Kir6.1) of the four major functional groups of Kir channels (Hibino et al., 2010). The alignment in Fig. 3 supplement shows that except for the selectivity filter, the viral channel has essentially no similarity with canonical Kir channels. Most importantly the viral channel contains none of the AAs, which are crucial for the function of canonical Kir channels, i.e., the AAs, which determine the steepness of rectification or polyamine sensitivity.

After discovering that inward rectification of the viral channel differs from Kir channels and that rectification is an inherent property of Kmpv_SP1_, we created a chimera with Kmpv_1_ to identify domains that might be responsible for rectification. First, we addressed the question of whether the functional differences in gating could be caused by the minor AA variations in the selectivity filter. Notably the inward rectifying virus channels do not include any of the specific AAs, which are typical for the filter region of Kir channels (Fig. 3 supplement). Neither the conserved disulfide bridge nor the pair of charged AAs, which are known to stabilize the filter of Kir channels (Yang et al., 1997; Leyland et al., 1999; Cho et al., 2000), are present in Kmpv_SP1_.

To examine the role of the filter in rectification we substituted the domain between the end of TM1 and the beginning of TM2 in Kmpv_SP1_, with the equivalent domain from Kmpv_1_ (ChA, Fig. 6A). When this chimera was expressed in HEK293 cells it exhibited the inward rectification of the parental channel that donated the TM domains (Fig. 6B,D).

**Figure 6:**
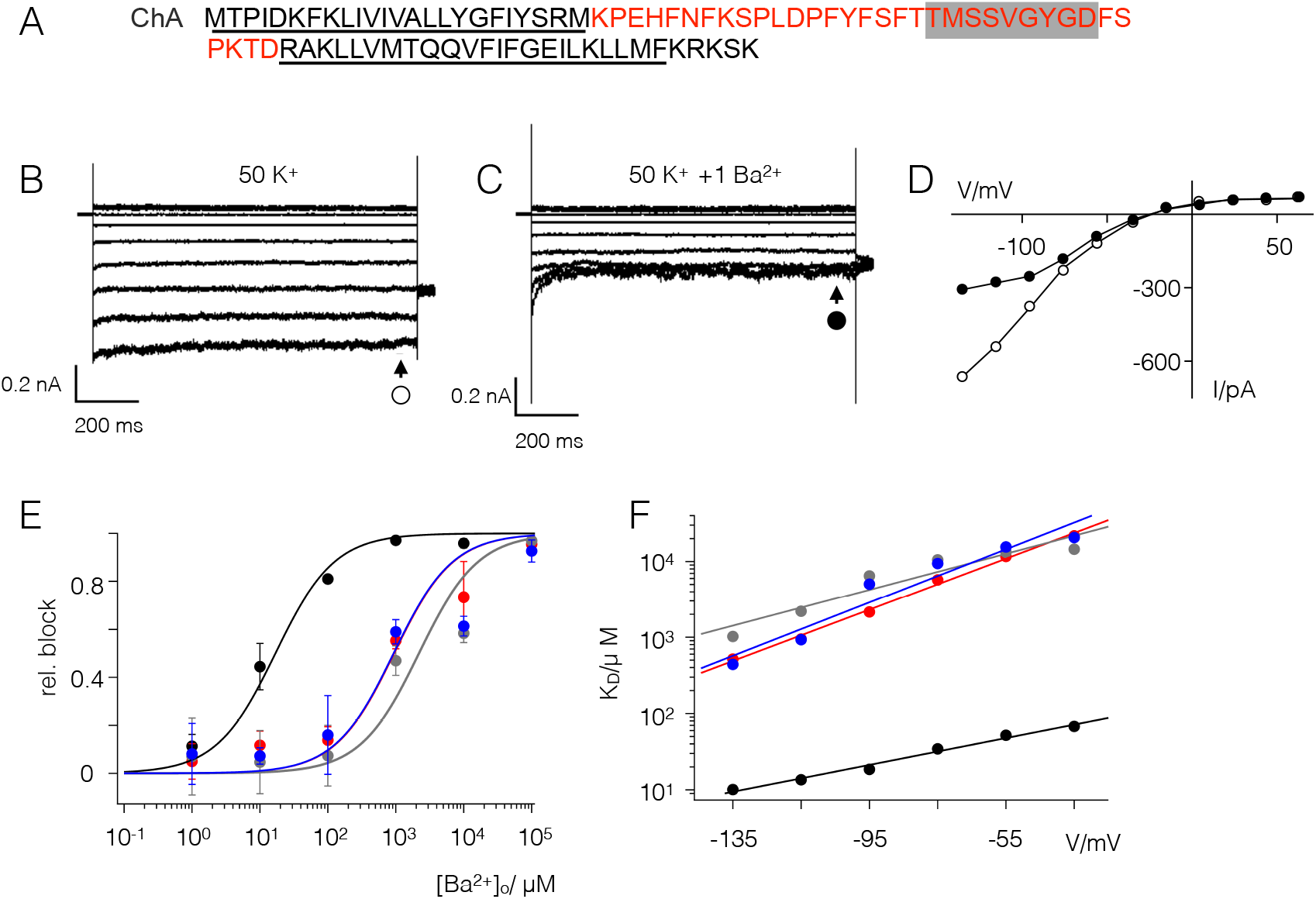
The filter domain of Kmpv_1_ is responsible for low Ba^2+^ sensitivity. **(A)** Chimera (ChA) comprising the filter of Kmpv_1_ (red) and the TM domains of Kmpv_SP1_ (black). The selectivity filter sequence is highlighted in grey; the estimated transmembrane domains are underlined. Exemplar currents as in Fig. 1 from HEK293 cells expressing ChA in bath solution with 50 mM K^+^ in the absence **(B)** and presence of 1 mM BaCl_2_ **(C)**. The corresponding I/V relations **(D)** show steady state currents in the absence (open circles) and presence (closed circles) of 1 mM Ba^2+^. **(E)** Relative block of Kmpv1 (black), Kmpv_SP1_ (red), chimera ChA (grey) and Kmpv_SP1_ mutant S53F (blue) at -115 mV as a function of extracellular Ba^2+^ concentration. Data (mean ± sd; n≥4) were fitted with the Hill equation. The K_D_ values for the Ba^2+^ block of both channels obtained from fits as in C are plotted as a function of voltage **(F)**. The K_D_ value increases ten fold per 48 mV (Kmpv_SP1_-S53F, blue) and 85 mV (ChA, grey) of negative voltage. For comparison the corresponding data of the wt channels are re-plotted.

To quantitatively compare the rectification properties of the wild type (wt) channel and the chimera we calculated the rectification index (R.I.) as the ratio of the chord conductance at +25 mV (G_25_) and -75 mV (G_−75_) according to Ozawa et al. (1991) with equation 2

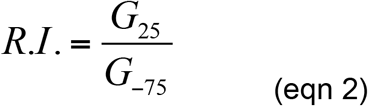

A comparison of the R.I. value between wt channel and chimera ChA shows that both channels exhibit no apparent difference in their voltage dependency (Fig. 7). This suggests that the filter domain is not the origin of rectification.

Further experiments showed that the chimera ChA maintained the low Ba^2+^ sensitivity of the parental filter domain (Fig. 6C-F). A scrutiny of the sequence of the two channels highlights a single AA difference in the vicinity of the selectivity filters with a Ser in Kmpv_SP1_ and a Phe in Kmpv_1_ (Fig. 1A). To test if this Ser is crucial for the high Ba^2+^ sensitivity of Kmpv_SP1_ it was mutated into a Phe. Functional testing of this mutant showed that the protein retained inward rectification but had lost its high sensitivity to Ba^2+^ (Fig. 6E,F); the dose response curve became similar to that of Kmpv_1_. The results of these experiments show that Ba^2+^ sensitivity is a function of the selectivity filter domain and that this property is not causally related to the mechanism of inward rectification.

**Figure 7:**
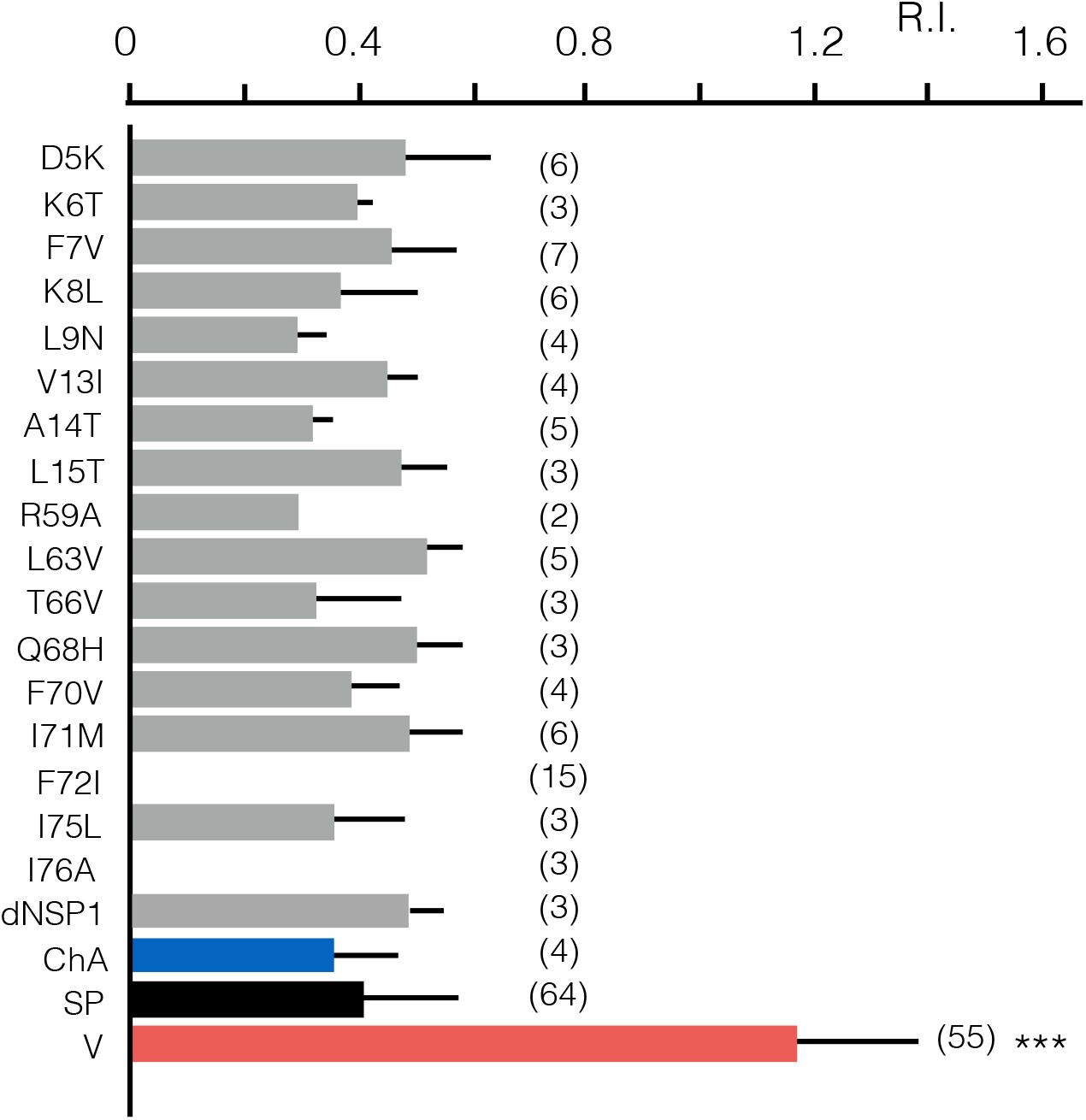
Rectification properties of Kmpv_1_, Kmpv_SP1_, chimera and point mutants. Rectification index (R.I.) of Kmpv_1_ (SP, red) Kmpv_SP1_ (V, black), chimera with Kmpv_1_ pore and Kmpv_SP1_ TM domains (ChA, blue) as well as mutants of Kmpv_SP1_ (grey). The latter comprise AAs in which Kmpv_1_ differs from the inward rectifiers Kmpv_PL1_ and Kmpv_SP1_ (see Fig. 1). The relevant AAs were mutated individually in Kmpv_SP1_ to match the sequence of the ohmic Kmpv1 channel. In the mutant dNSP1 three AAs (T2-I3) in the N-terminus of Kmpv_SP1_ were deleted. All constructs were expressed in HEK293 cells as in Figs 1 and 2. For mutant channels, which conducted significantly more inward current than un-transfected cells, the rectification index R.I. was calculated from eqn. 2. The mean R.I. values (± sd; number of recordings in brackets) are plotted. The R.I. value for Kmpv_1_ (V) is significantly (P<0.001; ***) larger than that of all other constructs. The R.I. value of Kmpv_SP1_ (SP) is not significantly different from all mutants/chimera; none of the mutations converted a rectifier into an ohmic conductor.

The results so far suggest that the TM segments rather than the filter region confer inward rectification. To test this hypothesis we identified the AAs, which are different in the N-and C-termini between the ohmic Kmpv_1_ channel on one side and the two rectifiers Kmpv_SP1_ and Kmpv_PL1_ on the other side (Fig. 1). The relevant AAs in Kmpv_SP1_ were mutated to those of Kmpv_1_ and the mutants expressed in HEK293 cells to search for a loss of inward rectification. All mutants except for F72I and L76A generated a functional channel in which the mean current significantly exceeded the mean current of untransfected cells; the R.I. values of all the functional mutants show that they all maintained an inward rectification (Fig. 7). The results of these experiments suggest that inward rectification is not conferred by a singular AA. Instead it seems to require tertiary or quaternary structural interactions of the transmembrane domains.

## Discussion

The present study shows that small viral K^+^ channels from environmental samples provide a rich protein library, which promotes understanding of basic structure/function correlates in K^+^ channels. Because of high evolutionary pressure on viral genomes these proteins exhibit an unbiased structural and functional diversity (Gazzarrini et al., 2004; Kang et al., 2004; Siotto et al., 2014; Rauh et al., 2017). The main outcome of the present comparative analysis of three viral channel proteins is that one is a simple ohmic conductor while the other two represent inward rectifiers. The latter resemble the function of Kir-type channels (Hibino et al., 2010). Like their larger relatives from bacteria and animals they conduct no substantial outward current at membrane potentials positive to the K^+^ equilibrium voltage, they are highly selective for K^+^ and undergo a voltage-dependent block by Ba^2+^. In spite of their functional similarities to strong rectifying Kir channels the mechanisms of gating in the viral channels must be fundamentally different. Unlike Kir channels the viral channels exhibit strong inward rectification without contributions from any cytosolic domains; the latter are important determinants of rectification in Kir channels (Baronas and Kurata, 2014). More importantly, while rectification in Kir channels is achieved by a voltage dependent block of the pore by cytosolic Mg^2+^ and polyamines, the viral channels rectify even in a pure KCl solution. Since the control experiments exclude contaminations in the solution as a source of channel blockers, we must assume that rectification is an inherent property of the viral channels. This interpretation is in line with previous reports, which had already shown that Kir mutants can acquire an intrinsic inward rectification. In the case of the ROMK1 channel it was found that the N171D mutation converted the channel into a strong inward rectifier even in the absence of Mg^2+^ (Lu and MacKinnon, 1994). This electrostatic effect however cannot explain the mechanism of rectification in the viral channels. The equivalent of an anionic AA equivalent to residue 171D in the ROMK1 mutant is not present in the viral channels (Fig. 3 supplement). If we consider that the alignment between Kir channels and Kmpv_SP1_ may not be correct the residue E74 of Kmpv_SP1_ could be in the same position of D171 in the ROMK1 mutant. This however provides no explanation for rectification in the viral channels since this Glu is also conserved in the Ohmic conducting Kmpv_1_ channel (Fig. 1).

A Kir channel with an intrinsic rectification was also found in response to the E224G mutant in Kir2.1 (Chang et al. 2007). In this case it was proposed that charges around the cytoplasmic pore may generate a local electrostatic potential at the channel entry for rectification. Since this domain is not present in the viral K^+^ channels (Fig. 3 supplement) it cannot account for their inherent rectification properties.

In the case of the intrinsic inward rectification of the viral channels we cannot explain the molecular mechanism of this voltage dependency. The finding that the pore loops of rectifying and non-rectifying channels can be swapped without any effect on rectification indicates that the pore domains with the selectivity filter architecture is not crucial for this voltage dependent phenomenon. This interpretation is supported by a mutation close to the selectivity filter, which alters the Ba^2+^ sensitivity of the inward rectifier. A reduction of Ba^2+^ sensitivity by nearly two orders of magnitude has in this case no impact on the rectifying properties of the channel. Hence the selectivity filter or the fine-tuning of the filter geometry seems to be irrelevant for an inherent inward rectification. This finding is unexpected considering the fact that inherent rectification is voltage and K^+^ sensitive and that the filter is the domain, which provides the selectivity (Doyle et al., 1998) for the channel and where the majority of the voltage drop in a K^+^ channel occurs (Contreras, et al., 2010; Andersson et al., 2018).

In the context of the differential Ba^2+^ sensitivity of the channels it is interesting to mention, that several previous structural and functional studies have identified the 4^th^ binding site, S4, in the filter of K^+^ channels as the primary site of the Ba^2+^ block (Chatelain et al., 2005; Piasta et al., 2011; Guo et al., 2014). The importance of S4 for the Ba^2+^ block was confirmed by experiments in which a Thr in this site, which is part of the consensus sequence of the filter, was mutated to a Ser. This mutation caused a drastic decrease in the sensitivity of another viral K^+^ channel but also in Kir channels to the Ba^2+^ block (Chatelain et al., 2005, 2009). In the three channels studied here the corresponding AA is already a Ser, which does not prevent a Ba^2+^ block with μM affinity in Kmpv_SP1_. Hence a Ser in this position is not solely responsible for a low Ba^2+^ affinity. As a main structural determinant of Ba^2+^ sensitivity the present data identify an AA in Kmpv_SP1_, which is on the opposite side of the selectivity filter that is exposed to the external medium. A single point mutation S53F is sufficient to shift the high Ba^2+^ sensitivity of Kmpv_SP1_ into that of Kmpv_1_. At this point we cannot answer the question of whether these data indicate an additional binding site on the extracellular side of the pore for Ba^2+^ or whether the mutation S53F affects the entry of Ba^2+^ into the filter and hence into its binding site deep in the electrical field. It is interesting to mention that an interplay of two residues on both sides of the selectivity filter is proposed for the Ba^2+^ block in the Kir2.1 channel (Alagem et al., 2001).

While the experiments exclude the filter as an important structure for inherent inward rectification in viral channels, they underscore the importance of the TM domains. The data also indicate that the conversion between a rectifying and ohmic conductance is not a matter of a single AA. The effective switch in voltage dependency, which was achieved by the exchange of the TM domains could neither be mimicked by the deletion of the first 3 AAs, which are lacking in Kmpv_1_ nor by point mutations of the AAs, which are different between the respective channels. Taken together these data suggest that the fold of the TM is crucial for generating an inherent inward rectification in Kmpv_SP1_. The question on how this is achieved and where the significant voltage dependency comes from remains unanswered.

## ACKNOWLEDGEMENTS

This research was supported by European Research Council (ERC) 2015 Advanced Grant 495 (AdG) n. 695078 noMAGIC (A.M., G.T.) the LOEWE initiative iNAPO (GT), Fondazione CARIPLO grant 2014-0796, by MIUR PRIN (Programmi di Ricerca di Rilevante Interesse Nazionale) 494 2015, (2015795S5W) to A.M and by the National Science Foundation grant no. 1736030 (JVE). We thank Dr. Fenja Siotto (Darmstadt) for initial cloning of the channel constructs.

## AUTHOR CONTRIBUTIONS

D.E, T.S and J.S. performed experiments, analyzed data and wrote parts of the paper; I.S., O.R. B.H. and I.S analyzed data. J.VE, A.M. and G.T. designed the work and wrote the paper.

## REFERENCES

Alagem, N., M. Dvir, and E. Reuveny. 2001. Mechanism of Ba^2+^-block of a mouse inwardly rectifying K^+^ channel: differential contribution by two discrete residues. J. Physiol. 534: 381–393.

Andersson, A. E. V., M.A. Kasimova, and L. Delemotte. 2018. Exploring the viral channel Kcv PBCV-1 function via computation. J. Membr. Biol. 251: 419–430.

H, Barry. P. 1994. JPCalc, a software package for calculating liquid junction potential corrections in patch-clamp, intracellular, epithelial and bilayer measurements and for correcting junction potential measurements. J. Neurosci. Meth. 51: 107–116.

Baronas, V. A., and H.T. Kurata. 2014. Inward rectifiers and their regulation by endogenous polyamines. Front. Physiol. 5: 325

Braun, C., T. Baer, A. Moroni, and G. Thiel. 2014. Pseudo painting/air bubble technique for planar lipid bilayers. J. Neurosci. Meth. 233: 13–17.

Chang, H.-K., S.-H. Yeh, and R.-C. Shieh. 2007. Charges in the cytoplasmic pore control intrinsic inward rectification and single-channel properties in Kir1.1 and Kir2.1 channels. J. Membr. Biol. 215: 181–193.

Chatelain, F. C., et al. 2005. The pore helix dipole has a minor mole in inward rectifier channel function. Neuron 47: 833–843.

Chatelain, F. C., et al. 2009. Selection of inhibitor-resistant viral potassium channels identifies a selectivity filter site that affects barium and amantadine block. PLoS One 4: e7496

Cho, H. C., R.G. Tsushima, T.T. Nguyen, H.R. Guy, and P.H. Backx. 2000. Two critical cysteine residues implicated in disulfide bond formation and proper folding of Kir2.1. Biochem. 39: 4649–4657.

Contreras, J. E. et al. 2010. Voltage profile along the permeation pathway of an open channel. Biophys. J. 99: 2863–2869.

Doyle, D. A., et al. 1998. The structure of the potassium channel: molecular basis of K^+^ conduction and selectivity. Science 280: 69–77.

Fujiwara, Y., and Y. Kubo. 2006. Functional roles of charged amino acid residues on the wall of the cytoplasmic pore of Kir2.1. J. Gen. Physiol. 127: 401–419.

Gazzarrini, S., et al. 2003. The viral potassium channel Kcv: structural and functional features. FEBS Lett. 552: 12–16.

Gazzarrini, S., et al. 2004. Long distance interactions within the potassium channel pore are revealed by molecular diversity of viral proteins. J. Biol. Chem. 279: 28443–28449.

Guo D., and Z. Lu. 2002. IRK1 inward rectifier K^+^ channels exhibit no intrinsic rectification. J. Gen. Physiol. 120: 539–551.

Guo, R., et al. 2014. Ionic interactions of Ba^2+^ blockades in the MthK K^+^ channel. J. Gen. Physiol. 144: 193–200.

Hamacher, K., T. Greiner, J.L. Van Etten, M. Gebhardt, C. Cosentino, A. Moroni, and G. Thiel. 2012. Phycodnavirus potassium ion channel proteins question the virus molecular piracy hypothesis. PLoSOne 7: e38826.

Hibino, H., et al. 2010. Inwardly rectifying potassium channels: their structure, function, and physiological roles. Physiol. Rev. 90: 291–366.

Huang CL, S. Feng, and D.W. Hilgemann. 1998. Direct activation of inward rectifier potassium channels by PIP_2_ and its stabilization by Gβγ. Nature 391: 803–806.

Kang, M., et al. 2004. Small potassium ion channel proteins encoded by chlorella viruses. Proc. Natl. Acad. Sci. U.S.A. 101: 5318–5324.

Kurata, H. T., L.R. Phillips, T. Rose, G. Loussouarn, S. Herlitze, H. Fritzenschaft, D. Enkvetchakul, C.G. Nichols, and T. Baukrowitz. 2004. Molecular basis of inward rectification: polyamine interaction sites located by combined channel and ligand mutagenesis. J. Gen. Physiol. 124: 541–554.

Kurata, H.T., W.W.L. Cheng, and C.G. Nichols. 2011. Polyamine block of inwardly rectifying potassium channels. Meth. Mol. Biol. 720: 113–126.

Kuang, Q., et al. 2015. Structure of potassium channels. Cell. Mol. Life Sci. 72: 3677–3693.

Leyland, M. L., C. Dart, P. Spencer, M. J. Sutcliffe, and P.P. Stanfield. 1999. The possible role of a disulphide bond in forming functional Kir2.1 potassium channels. Pflügers Arch.-Europ. J. Physiol. 438: 778–781.

Lopatin, A.N., E.N. Makhina, and C.G. Nichols. 1994. Potassium channel block by cytoplasmic polyamines as the mechanism of intrinsic rectification. Nature 372: 366–369.

Lopatin, A. N., and C. G. Nichols. 1996. K^+^ dependence of open-channel conductance in cloned inward rectifier potassium channels (IRK1, Kir2.1). Biophys. J. 71: 682–694.

Lu, Z., and R. MacKinnon. 1994. Electrostatic tuning of Mg^2+^ affinity in an inward-rectifying K^+^ channel. Nature 371:243l246.

Lu, Z. 2004. Mechanism of rectification in inward-rectifying K^+^ channels. Annu. Rev. Physiol. 66: 103–129

MacKinnon, R., 2003. Potassium channels. FEBS Lett. 555: 62–65.

Makhina, E. N., et al. 1994. Cloning and expression of a novel human brain inward rectifier potassium channel. J. Biol. Chem. 269: 20468–20474.

Minor, D.L. 2009. Searching for interesting channels: pairing selection and molecular evolution methods to study ion channel structure and function. Mol. Biosyst. 5: 802–810.

Nichols, C.G., and S.J. Lee. 2018. Polyamines and potassium channels: A 25-year romance. J. Biol. Chem. 293: 18779–18788.

Ozawa, S., et al. 1991. Two types of kainate response in cultured rat hippocampal neurons. J. Neurophysiol. 66: 2–11.

Perier, F., et al. 1994. Primary structure and characterization of a small-conductance inwardly rectifying potassium channel from human hippocampus. Proc. Natl. Acad. Sci. U.S.A. 91: 6240–6244.

Perozo, E. 2002. New structural perspectives on K^+^ channel gating. Structure 10: 1027–1029.

Piasta, K. N., et al. 2011. Potassium-selective block of barium permeation through single KcsA channels. J. Gen. Physiol. 138: 421–436.

Rauh, O., et al. 2017. Identification of intrahelical bifurcated H-bonds as a new type of gate in K^+^ channels. J. Am. Chem. Soc. 139: 7494–7503.

Roux, B. 2017. Ion channel and ion selectivity. Essays Biochem. 61:201–209.

Siotto, F., et al. 2014. Viruses infecting marine picoplancton encode functional potassium ion channels. Virology 466–467: 103–111.

Spassova, M., and Z. Lu. 1999. Tuning the voltage dependence of tetraethylammonium block with permeant ions in an inward-rectifier K^+^ channel. J. Gen. Physiol. 114:415–26

Thiel, G., A. Moroni, D. Dunigan, and J.L. Van Etten. 2010. Initial events associated with virus PBCV-1 infection of *Chlorella* NC64A. Prog. Bot. 71: 169–183.

Thiel, G., et al. 2011. Minimal art: or why small viral K^+^ channels are good tools for understanding basic structure and function relations. Biochim. Biophys. Acta. 1808: 580–588.

Winterstein, L.-M., et al. 2018. Reconstitution and functional characterization of ion channels from nanodiscs in lipid bilayers. Reconstitution and functional characterization of ion channels from nanodiscs in lipid bilayers. J. Gen Physiol. 150: 637–646.

Yang, J., M. Yu, Y.N. Jan, and L.Y. Jan. 1997. Stabilization of ion selectivity filter by pore loop ion pairs in an inwardly rectifying potassium channel. Proc. Natl. Acad. Sci. U.S.A. 94: 1568–1572.

