## supplement figures for "A small viral potassium ion channel with an inherent inward rectification"

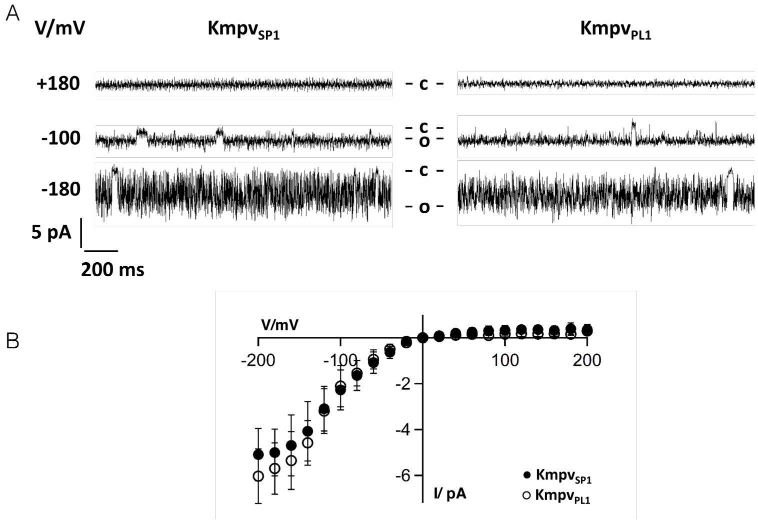


**Figure 1 supplement: Kmpv$_{PL1}$ has the same inward rectification as Kmp$_{SP1}$. (A)** Exemplar current traces of Kmpv$_{PL1}$ and Kmpv$_{SP1}$ activity over a range of clamp voltages in symmetrical KCl buffer (100 mM KCl + 10 mM HEPES, pH 7). Both channel proteins were translated *in vitro* as in Fig. 4. The Kmpv$_{PL1}$ channel exhibits the same high open probability with flicker type fluctuations as Kmpv$_{SP1}$. **(B)** Mean I/V relations of average currents (mean ± sd; n≥9) of both channels measured over clamp steps of 60 sec.

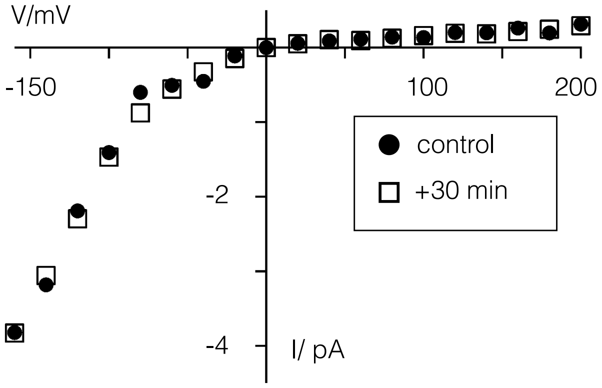


**Figure 2 supplement: Long term incubation of Kmpv$_{SP1}$ in pure KCl buffer does not compromise rectification.** I/V relation of mean currents measured in planar lipid bilayer with Kmpv$_{SP1}$ protein immediately after functional reconstitution (circles) and 30 min later (squares). Data were recorded as in Fig. 4.

```
KmpvSP1  ------------------------------------------------------------
Kir1.1   MNASSRNVFDTLIRVLTESMFKHLRKWVVTRFFGHSRQ--------RARLVSKDGRCNIE
Kir6.1   MLARKSIIPEEYVLARIAAENLRKP-----------RIRDRLP---KARFIAKSGACNLA
Kir2.1   MGSVRT---NRYSIVSSEEDGMKLATMAVANGFGNGKSKVRTRQQCRSRFVKKDGHCNVQ
Kir3.1   MSALRRKFGDDYQVVTTSSSGSGLQ----PQGPGQDPQQQLVPKKKRQRFVDKNGRCNVQ

KmpvSP1  ------------------MTPID---KFKLIVIV-----------ALLYGFIYSRMDPEE
Kir1.1   FGNVEAQS-RFIFFVDIWTTVLDLKWRYKMTIFITAFLGSWFFFGLLWYAVAYIHKDLPE
Kir6.1   HKNIREQG-RFL--QDIFTTLVDLKWRHTLVIFTMSFLCSWLLFAIMWWLVAFAHGDIYA
Kir2.1   FINVGEKGQRYL--ADIFTTCVDIRWRWMLVIFCLAFVLSWLFFGCVLWLIALLHGDLDA
Kir3.1   HGNLGSETSRYL--SDLFTTLVDLKWRWNLFIFILTYTVAWLFMASMWWVIAYTRGDLNK
                            * :*   .  : ::             : :  .   . *

KmpvSP1  F------------------GFSSPLDPYYFSFTTMSSVGYGDS--SPKTDRAKLLVMTQQ
Kir1.1   F-------HPSANHTPCVENINGLTSAFLFSLETQVTIGYGFRCVTEQCATAIFLLIFQS
Kir6.1   YMEKSGMEKSGLESTVCVTNVRSFTSAFLFSIEVQVTIGFGGRMMTEECPLAITVLILQN
Kir2.1   S-------KEG---KACVSEVNSFTAAFLFSIETQTTIGYGFRCVTDECPIAVFMVVFQS
Kir3.1   A-------HVG-NYTPCVANVYNFPSAFLFFIETEATIGYGYRYITDKCPEGIILFLFQS
                            . .   .: * : .  ::*:*     : :   .   :.:*.

KmpvSP1  V-------FIGGEIL-KLLMFKRKSK
Kir1.1   ILGVIINSFMCGAILAKISRPKKRAKTITFSKNAVISKRGGKLCLLIRVANLRKSLLIGS
Kir6.1   IVGLIINAVMLGCIFMKTAQAHRRAETLIFSRHAVIAVRNGKLCFMFRVGDLRKSMIISA
Kir2.1   IVGCIIDAFIIGAVMAKMAKPKKRNETLVFSHNAVIAMRDGKLCLMWRVGNLRKSHLVEA
Kir3.1   ILGSIVDAFLIGCMFIKMSQPKKRAETLMFSEHAVISMRDGKLTLMFRVGNLRNSHMVSA
         :       .: * :: *    :..   * : : :: :..*           ..:: :  :.

KmpvSP1  -----------------------------------------------------------
Kir1.1   HIYGKLLKTTVTPEGETIILDQININFVVDAGNENLFFISPLTIYHVIDHNSPFFHMAAE
Kir6.1   SVRIQVVKKTTTPEGEVVPIHQLDIPVDNPIESNNIFLVAPLIICHVIDKRSPLYDISAT
Kir2.1   HVRAQLLKSRITSEGEYIPLDQIDINVGFDSGIDRIFLVSPITIVHEIDEDSPLYDLSKQ
Kir3.1   QIRCKLLKSRQTPEGEFLPLDQLELDVGFSTGADQLFLVSPLTICHVIDAKSPFYDLSQR

KmpvSP1  ------------------------------------------------------------
Kir1.1   TLLQQDFELVVFLDGTVESTSATCQVRTSYVPEEVLWGYRFAPIVSKTKEGKYRVDFHNF
Kir6.1   DLANQDLEVIVILEGVVETTGITTQARTSYIAEEIQWGHRFVSIVTE-EEGVYSVDYSKF
Kir2.1   DIDNADFEIVVILEGMVEATAMTTQCRSSYLANEILWGHRYEPVLFE-EKHYYKVDYSRF
Kir3.1   SMQTEQFEIVVILEGIVETTGMTCQARTSYTEDEVLWGHRFFPVISL-EEGFFKVDYSQF

KmpvSP1  ------------------------------------------------------------
Kir1.1   SKTVEVETPH--------------------------------------------------
Kir6.1   GNTVKVAAPR--------------------------------------------------
Kir2.1   HKTYEVPNTP--------------------------------------------------
Kir3.1   HATFEVPTPPYSVKEQEEMLLMSSPLIAPAITNSKERHNSVECLDGLDDITTKLPSKLQK

KmpvSP1  ------------------------------------------------------------
Kir1.1   ------------C-----------------------AMCLYNEKDVRARMKRGYDNP---
Kir6.1   ------------CSARELDEKPSILIQTLQKSELSHQNSLRKRNSMRRNNSMRRNNSIRR
Kir2.1   -----------LCSARDLAEKKYILSN-------ANSFCYENEVALTSKEEDDSENGVPE
Kir3.1   ITGREDFPKKLLRMSSTTSEKAYSLGDLPMKLQRISSVPGNSEEKLVSKTTKMLSDPMSQ

KmpvSP1  -------------------------------------
Kir1.1   ----------------------NFILSEVNETDDTKM
Kir6.1   NNSSLMVPKVQ-------------FMTPEGNQNTSES
Kir2.1   STSTDTPPDID------LHNQASVPLEPRPLRRESEI
Kir3.1   SVA-DLPPKLQKMAGGAARMEGNLPAKLRKMNSDRFT
```

**Figure 3 supplement: Viral inward rectifier shares no structural similarity with canonical Kir channels.** Alignment of $Kmpv_{SP1}$ with canonical Kir channels. The four major functional subgroups (Hibino et al., 2010) of Kir channels are represented by the human Kir1.1 (BAG36779.1), Kir2.1 (NP_000211), Kir3.1 (P48549) and Kir6.1 (Q15842.1). Alignment performed with MUSCLE algorithm (https://www.ebi.ac.uk/Tools/msa/muscle/). Kir typical AAs are highlighted in the Kir2.1 sequence including: i) two Cys residues and charged AAs in the filter (in blue; Baronas and Kurata 2014), ii) the critical AA, which determines between weak and strong rectification (in green; Hibero et al., 2010; Baronas and Kurata, 2014), iii) residues which induce strong rectification when an acidic substitution is introduced in Kir6.2 (in turquoise; Baronas and Kurata, 2014; Nichols and Lee, 2018), (iv) AAs for $PIP_2$ interaction (in red; Hibero et al. 2010), (v) charged AAs on the wall of the cytoplasmic pore (in yellow; Fujiwara and Kubo 2006). Asterisks (*) indicate conserved residues, colons (:) indicate residues with strongly similar properties, and periods (.) indicate residues with weakly similar properties.
